# Functional variations in gamma-secretase activity are critical determinants of the clinical, biomarker, and cognitive progression of autosomal dominant Alzheimer’s disease

**DOI:** 10.1101/2023.07.04.547688

**Authors:** Stephanie A. Schultz, Lei Liu, Aaron P. Schultz, Colleen D. Fitzpatrick, Raina Levin, Jean-Pierre Bellier, Zahra Shirzadi, Nelly Joseph-Mathurin, Charles D. Chen, Tammie L.S. Benzinger, Gregory S. Day, Martin R. Farlow, Brian A. Gordon, Jason J. Hassenstab, Clifford R. Jack, Mathias Jucker, Celeste M. Karch, Jae-Hong Lee, Johannes Levin, Richard J. Perrin, Peter R Schofield, Chengjie Xiong, Keith A. Johnson, Eric McDade, Randall J. Bateman, Reisa A. Sperling, Dennis J. Selkoe, Jasmeer P. Chhatwal, Dominantly Inherited Alzheimer Network Investigators

## Abstract

**Background:** The balance between production, clearance, and toxicity of Aβ peptides is central to Alzheimer’s disease (AD) pathobiology. Though highly variable in terms of age at symptom onset (AAO), hundreds of variants in *PSEN1* cause autosomal dominant forms of AD (ADAD) with nearly complete penetrance. PSEN1 forms the catalytic core of the γ-secretase complex and thereby directly mediates the production of longer, aggregation-prone Aβ peptides relative to shorter, non-aggregating peptides. We hypothesized that the broad AAO and biomarker heterogeneity seen across ADAD would be predictable based on mutation-specific differences in the production of Aβ species.

**Methods:** Aβ-37, 38, 40, 42, and 43 production was quantified from 161 unique *PSEN1* variants expressed in HEK293 cells. Prediction of AAO was carried out in 106 variants with available AAO and then replicated in 55 variants represented across 190 PSEN1 mutation carriers who have detailed cognitive and biomarker data from the Dominantly Inherited Alzheimer’s Network (DIAN).

**Results:** Variations in Aβ production across the 161 mutations examined in cell-based models were highly predictive of AAO. In those with corresponding *in vivo* data from the DIAN study, our cell-based γ-secretase composite was strongly associated with biomarker and cognitive trajectories.

**Conclusions:** These findings elucidate the critical link between γ-secretase function, Aβ production, and AD progression and offer mechanistic support for the amyloid hypothesis. The approach used here represents a powerful tool to account for heterogeneity in disease progression in ADAD clinical trials and to assess the pathogenicity of variants of unknown significance or with limited family history.

## INTRODUCTION

An increase in longer, aggregation-prone Aβ fragments (e.g., Aβ 42 and 43) relative to shorter, non-aggregating fragments (e.g., Aβ 37 and 38) is a critical initiating pathogenic event in both late-onset, sporadic AD^1^ (LOAD) and Autosomal Dominant AD^2–4^ (ADAD). The balance between production of aggregating and non-aggregating forms of Aβ is a direct result of the efficiency and kinetics with which the γ-secretase proteolytic complex sequentially cleaves β-amyloid precursor protein^5–10^ (APP). Subtle alterations in γ-secretase function can lead to profound neurodegenerative and cognitive consequences, while modulation of γ-secretase has therapeutic potential in both LOAD and ADAD^8, 9, 11–13^. Over 200 pathogenic variants in *Presenilin-1* (*PSEN1*), the key catalytic subunit in the γ-secretase complex, have been identified and are the most common causes of ADAD^14^. Mirroring the heterogeneity in clinical and AD biomarker progression seen in LOAD, there is striking heterogeneity in age of symptom onset (AAO)^15–17^ and rates of cognitive and biomarker change^18–22^ across carriers of *PSEN1* pathogenic variants. We hypothesized that this heterogeneity may be strongly tied to underlying mutation-level variation in γ-secretase function.

Intriguingly, prior work suggests some pathogenic *PSEN1* variants may not significantly alter the absolute amounts of Aβ42 production^23, 24^, indicating that more thorough investigation is needed to understand how these pathogenic variants lead to ADAD and the mechanisms that cause earlier vs. later disease onset and progression. To this end, findings from our group^25^ and others^26^ suggest that probing the ratios of short-to-long Aβ peptide production across ADAD pathogenic variants may help us understand the observed heterogeneity in these populations. However, this initial work examined only a small number of variants, used only AAO information derived from the literature, and lacked biomarker, clinical, and cognitive data.

Here, we leverage a live-cell-based model system and newly developed immunoassays to functionally characterize APP processing by γ-secretase across a large set of *PSEN1* pathogenic variants with AAOs ranging from the 20s to the 70s. We then assess how this mutation-level characterization of γ-secretase function may explain heterogeneity in neuroimaging and biofluid biomarkers, as well as clinical measures among *PSEN1* carriers with available data from the Dominantly Inherited Alzheimer’s Network Observational Study (DIAN-Obs).

## METHODS

### Functional assays of γ-secretase function

161 unique *PSEN1* variants, including 55 unique variants represented in DIAN-Obs and data from 106 unique variants previously characterized in Liu et al., 2022,^25^ and wild-type (normal) *PSEN1* was characterized for Aβ peptide production and γ-secretase function. Using methods detailed in supplemental methods and a prior report using this model system, each *PSEN1* variant or normal *PSEN1* was sub-cloned and placed into the PcDNA 3.1 expression vector^25^. Subsequently, each variant was co-transfected with a plasmid encoding wild-type (normal) human APP-C99 into HEK293T cells genetically depleted of *PSEN1* and *PSEN2*^25^. Creation of this cell line and transient transfection procedures are described in supplemental methods. Conditioned media from transfected HEK cells was harvested and diluted with 1% BSA in wash buffer (TBS supplemented with 0.05% Tween). Aβ37, 38, 40, 42, and 43 immunoassays were performed in triplicate and averaged as previously described^25^ (see Figure S1 and supplemental methods). Absolute peptide production levels and relevant ratios were examined as percent relative to the production seen in the same model system following transfection with normal (wild-type) *PSEN1*.

### Participants

Participants provided written informed consent prior to the completion of any study procedures, in accordance with Institution Review Board policy at each participating site. Data were included from 190 *PSEN1* pathogenic variant carriers enrolled in DIAN-Obs, each carrying one of 55 unique *PSEN1* pathogenic variants characterized via cell-based methods (see Figure 1A, Table S1, and Table S2). Detailed protocols for DIAN have previously been published and are described in supplemental methods. For DIAN-Obs participants, the familial AAO was determined through structured interviews to determine the age of onset of progressive cognitive decline for the participant’s first degree relative(s)^16^.

**Figure 1.**
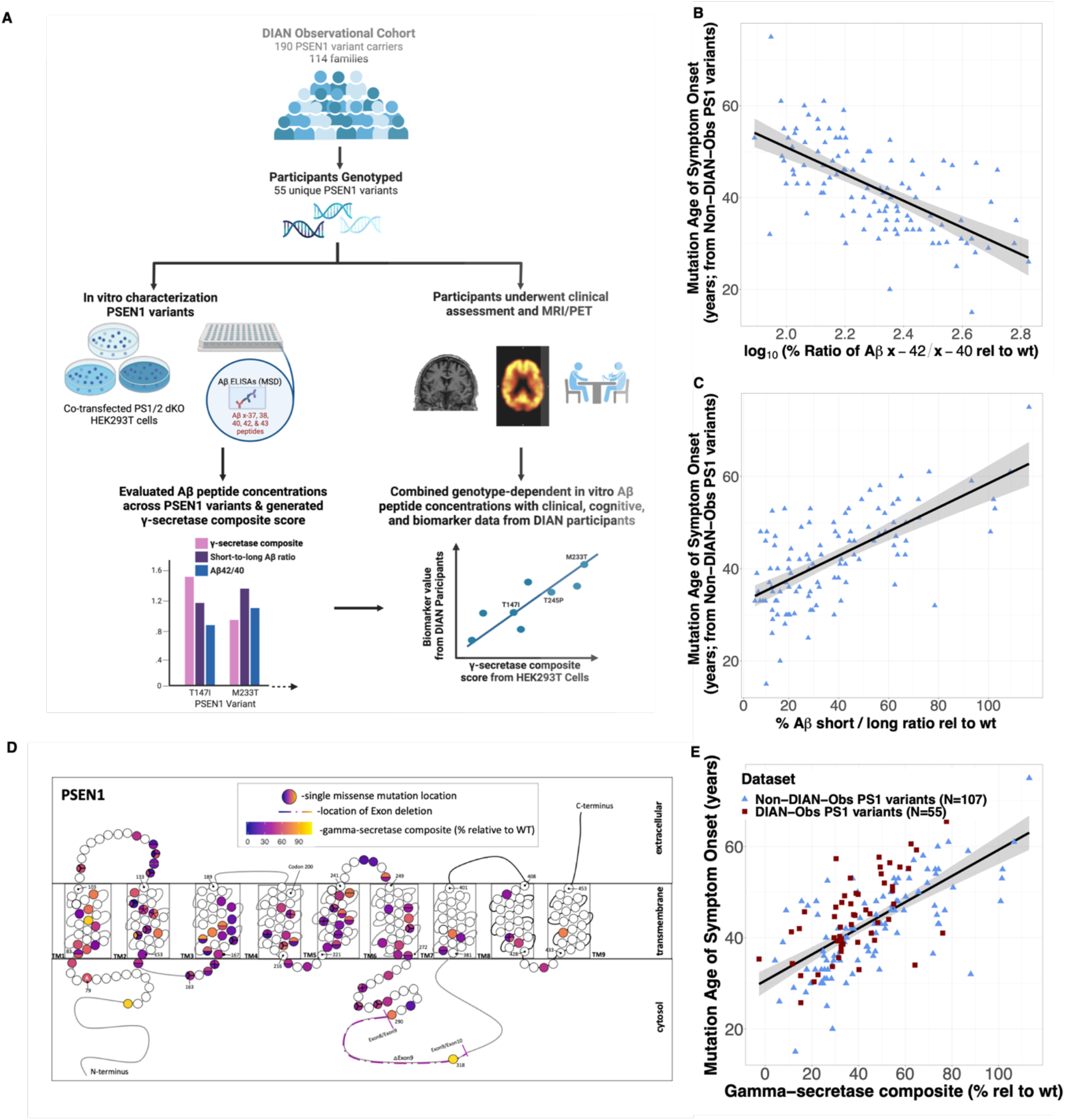
Cell-based measures of γ-secretase function are tightly associated with age of symptom onset across two cohorts representing 161 unique *PSEN1* variants. **(A)** Schematic summarizing the characterization of variant-level γ-secretase function and correlations with age of symptom onset, cognitive, and biomarker data for each variant (created with BioRender.com). γ-secretase function was summarized using two ratios of Aβ peptide production derived from cell models (**B**) Cell-based 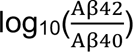 levels and (**C**) a ratio of short (fully γ-secretase processed) to long (incompletely γ-secretase processed) Aβ fragments 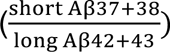. Both ratios were independently associated with age of symptom onset in an initial set of 106 *PSEN1* variants (B-C). To reduce comparisons going forward, both ratios were combined into a single composite measure (γ-secretase composite, expressed as a percentage of normal γ-secretase function; D). This composite measure correlated with age of symptom onset in both the initial set of 106 *PSEN1* variants and in a second set of 55 *PSEN1* variants represented in DIAN-Obs (E). Individual data points are jittered to maintain blinding.

Clinical evaluators were blind to the mutation status of participants. Participants underwent comprehensive clinical and cognitive evaluations. Clinical Dementia Rating (CDR®) Sum of Boxes (SB) was assessed for each participant using structured interviews, as previously described^27^. Mini-Mental State Examination (MMSE) and Wechsler Memory Scale-Revised Logical Memory Delayed Recall^28^ were selected a priori as our cognitive measures of interest. All DIAN-Obs participants with available Aβ positron emission tomography (PET), magnetic resonance imaging (MRI), and at least one cognitive assessment were included (Figure 1A; DIAN-Obs data freeze version 15; last data from June 30, 2020; see supplemental methods). A subset of these individuals also had [^18^F]Fluorodeoxyglucose (FDG)-PET (n=162) and cerebrospinal fluid (CSF) (n=157) available for analyses. Longitudinal data were available from 154 DIAN participants (Table S3). For the 106 *PSEN1* variants studies previously in Liu et al. (non-overlapping with the 55 variants from DIAN-Obs), AAO was derived from prior literature, as previously described^25^.

### Imaging and Fluid Biomarker Analyses

Protocols for Aβ and brain metabolism PET, MRI, and CSF studies are described in detail in supplemental methods and in previously published work^22, 27, 29–33^. Cerebral Aβ load was measured using [^11^C]Pittsburgh Compound B (PiB) PET, and brain glucose metabolism was measured with [^18^F]FDG-PET. Partial volume corrected values were used in all analyses. FreeSurfer v 5.3 defined regional measures were derived from MRI data. CSF Aβ42, Aβ40, and phosphorylated-tau181 levels were measured using an automated immunoassay system (LUMIPULSE G1200, Fujirebio, Malvern, PA). Phosphorylated-tau 217 (pT217/T217) values were derived from immunoprecipitation-mass spectrometry (IP-MS) as previously described^29^. See supplemental methods for additional details.

### Statistical Analyses

Analyses were conducted using R (version 4.0.3, R Foundation for Statistical Computing). Pearson correlations were used to assess associations of Aβ production and γ-secretase function with AAO. A summary measure of variant-level Aβ production and γ-secretase function (γ-secretase composite) was developed to reduce the number of statistical comparisons and facilitate comparisons between mutant and wild-type *PSEN1*, as described in the Results.

Linear mixed effect models including fixed effects for age, sex, and γ-secretase composite score were employed to assess associations between the cell-based γ-secretase function and multi-modal biomarker and clinical data from DIAN. Similar to prior work from DIAN, a random effect for family membership was included in each linear mixed effects model. Separate models were used to assess associations with each clinical, cognitive, or biomarker outcome measure, focusing on CSF Aβ 42/40, CSF phosphorylated-tau 181, CSF phosphorylated-tau 217, hippocampal volume, CDR-SB, MMSE, and logical memory delayed (see supplemental methods). Years of education was included as an additional fixed effect term in models examining cognitive and clinical outcomes. Longitudinal models were similarly constructed but additionally included interactions of each fixed effect with time and terms allowing both a random intercept and slope. Additional exploratory analyses made use of a tertile split of γ-secretase composite scores to visualize how clinical, cognitive, and biomarker trajectories can be understood in the context of γ-secretase function.

## RESULTS

### Cell-based measures of γ-secretase function are associated with age of symptom onset

Aβ 37, 38, 40, 42, and 43 production was assessed in 161 unique *PSEN1* variants, including 55 *PSEN1* variants represented in DIAN (Table S1) and 106 non-overlapping variants previously characterized in Liu et al., 2022^25^. Grouping together Aβ production for all 161 variants, we observed reductions in Aβ37, 38, and 40 compared to normal levels (median % reduction relative to normal = −56.8, −48.6, and −22.8%, respectively), suggesting that, as a group, *PSEN1* pathogenic variants decrease the processive γ-cleavage of Aβ by γ-secretase (γ-processivity), leading to a decrease in these fully processed (shorter) Aβ peptides. Additionally, relative levels of Aβ42 and 43 across variants were higher in *PSEN1* pathogenic variants compared to normal levels (median % increase relative to normal= 66.5% and 47.1%; respectively).

We first assessed relationships between the relative concentrations of Aβ species produced by each variant and AAO in a set of 106 variants not represented in the DIAN study. To limit comparisons with multiple ratios when assessing relationships between γ-secretase function and AAO, we leveraged prior literature^1, 25, 26, 34, 35^ suggesting the Aβ 42/40 ratio and a measure of the efficiency of successive cleavage (γ-processivity) of Aβ fragments can be used to summarize γ-secretase function. These two measures were operationalized as 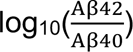 (a measure of the relative production of aggregation-prone Aβ) and a ratio of short (fully γ-secretase processed) to long (incompletely γ-secretase processed) Aβ fragments 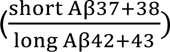. As expected, 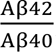, and 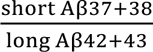 ratio levels across *PSEN1* pathogenic variants were both abnormal (decreased processivity) compared to normal levels (median % change relative to the normal = 108.9 and −71.3%). In the initial sample of 106 *PSEN1* pathogenic variants from Liu et al., both higher 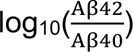 (r[105] = −0.64, p = 2.20e-13) and lower 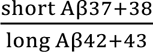 (r[105] = 0.66, p =2.28e-14) levels were associated with earlier AAO (Figure 1B-C). Both ratios remained significantly associated when entered together into a single model predicting AAO (Table S4), suggesting that they capture unique variance with respect to the pathogenicity of each variant. Accordingly, these two ratios were combined into a single summary measure of variant-level γ-secretase function (γ − secretase 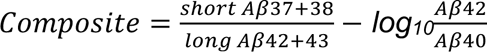). To facilitate visualization and comparison, the γ-secretase composite is expressed as a percentage of wild-type γ-secretase function.

In both the subset of variants represented in the DIAN-Obs and remaining variants characterized previously by Liu et al^25^, γ-secretase composite scores were lower in all pathogenic variants compared to normal (average of 60.4% reduction relative to normal). The γ-secretase composite had a the highest degree of correlation with AAO (Liu et al cohort: r[105] = 0.71, p <2.2e-16; DIAN-Obs cohort: r[53] = 0.61, p = 6.10e-07; Figure 1E), as compared to using individual Aβ peptide levels, 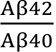 levels, or 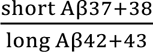 ratio levels in the initial set of 106 *PSEN1* variants and in the 55 variants represented in DIAN (see above). To demonstrate the potential utility of using the γ-secretase composite to predict an approximate AAO for novel pathogenic variants and/or when a clinical history is not available, we list the γ-secretase composite derived AAO (AAO_gsc_) values for the variants examined here (Table S1). Notably, γ-secretase composite values did not map in a straightforward manner onto the location of the corresponding pathogenic variant within *PSEN1*, and substantial variation within a single codon in terms of γ-secretase function was observed for several sites in which more than one variant was characterized (Table S1; Figure 1D).

### Lower cell-based γ-secretase composite scores are associated with more abnormal AD imaging and fluid biomarkers

Lower γ-secretase composite scores (decreased γ-secretase function relative to normal) were associated with higher levels of mean cortical Aβ burden (B[SE] = −0.03[0.01], p =4.06e-07; Figure 2A) as assessed by Aβ PET, after controlling for demographic factors. Exploratory regional analyses revealed widespread associations between *in vivo* Aβ burden and the cell-based γ-secretase composite, with the strongest associations seen in the precuneus, and anterior and posterior cingulate (Figure 2D).

**Figure 2.**
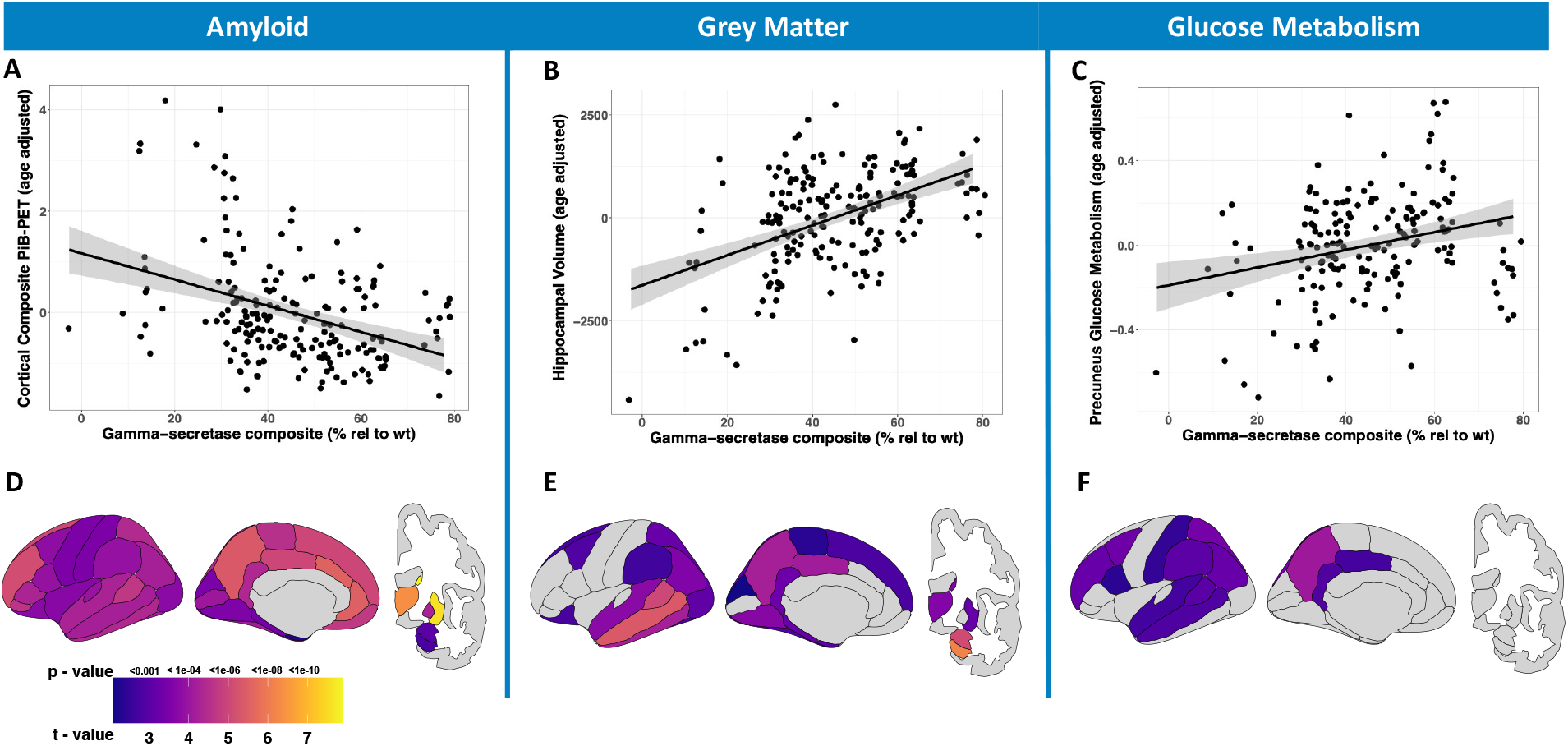
Lower γ-secretase function across variants is associated with more abnormal AD imaging biomarkers. Association between variant-level γ-secretase function and in *vivo* imaging biomarker data from carriers of corresponding *PSEN1* pathogenic variants is shown for cross-sectional Aβ PET (age-adjusted PiB-PET SUVR cortical composite, **A**), age- and intracranial volume-adjusted bilateral hippocampal grey matter volume derived from structural MRI (**B**), and age-adjusted bilateral precuneus FDG-PET SUVR (**C**) from 190 *PSEN1* pathogenic variant carriers (55 unique variants represented). Individual data points are jittered to maintain blinding of the data. Exploratory, regional analyses comparing PiB-PET SUVRs (**D**), grey matter volumes (**E**), and FDG-PET SUVRs (**F**) across a range of γ-secretase function, with colors indicating t- and p-values derived from the comparison of neuroimaging data from *PSEN1* carriers with variants in the lowest (most pathogenic) vs. highest (least pathogenic) tertiles of γ-secretase composite function. Models additionally adjusted for age and sex. Individual data points are jittered to maintain blinding.

We next examined associations between the variant-level γ-secretase composite and commonly used imaging biomarkers of neurodegeneration, including structural MRI and FDG-PET, in DIAN-Obs participants carrying one of the 55 of the *PSEN1* variants characterized *in vitro*. Both the MRI-based measures hippocampal volume (B[SE] = 44.22[6.4], p = 5.15e-10; Figure 2B) and precuneus FDG-PET signal (B[SE] = 0.006[0.001], p = 5.38e-05; Figure 2C) were highly associated with the γ-secretase composite, after controlling for demographic factors. Exploratory regional analyses revealed that the cell-based γ-secretase composite significantly accounted for widespread grey matter volume differences across *PSEN1* variant carriers, most notably in the hippocampus, amygdala, temporal regions, precuneus, and posterior cingulate (Figure 2E), as well as differences in glucose metabolism including the precuneus, parietal, and frontal regions (Figure 2F).

Using established CSF measures of AD pathology, we observed that lower variant-specific γ-secretase composite scores were associated with lower CSF Aβ 42/40 (B[SE] = 6.3e-04[1.5e-04], p = 4.46e-05; Figure 3A), higher CSF ELISA log_10_(phosphorylated-tauT181) levels (B[SE] = −0.009[0.002], p =4.47e-06; Figure 3B), and higher CSF IP-MS log_10_(pT217/T217) levels (B[SE] = −0.01[0.002], p =1.98e-05; Figure 3C).

**Figure 3.**
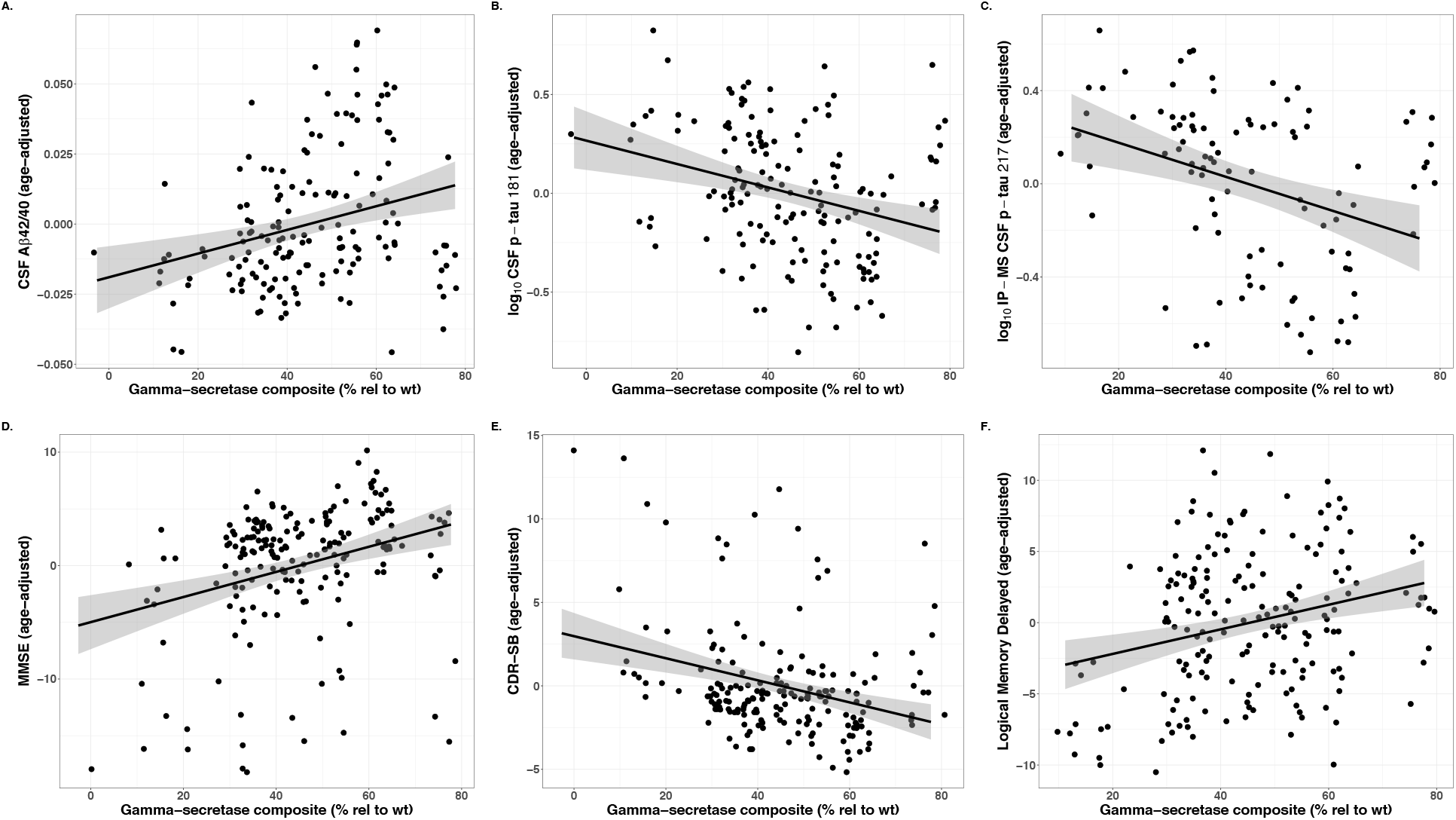
Lower γ-secretase function across variants is associated with more abnormal biofluid, clinical, and cognitive measures in ADAD. Associations between γ-secretase function for each *PSEN1* variant and cross-sectional CSF Aβ 42/40 (**A**), age-adjusted CSF log_10_ pT181 (**B**), CSF log_10_ pT217/T217 (**C**), age-adjusted MMSE (**D**), age-adjusted CDR-SB (**E**), and age-adjusted logical memory-delayed (**F**) values from 190 *PSEN1* pathogenic variant carriers (55 unique variants represented) are shown. Individual data points are jittered to maintain blinding. CSF = Cerebrospinal Fluid; pT = phosphorylated tau; MMSE = Mini Mental State Examination scores; CDR-SB = Clinical Dementia Rating Scale-Sum of Boxes.

### γ-secretase composite scores are associated with clinical and cognitive measures in *PSEN1* pathogenic variant carriers in DIAN-Obs

The cell-derived γ-secretase composite levels predicted changes of MMSE (B[SE] = 0.11[.03], p = 2.17e-04; Figure 3D), CDR-SB (B[SE] = −0.07[0.02], p = 4.86e-05; Figure 3E), and logical memory-delayed scores (B[SE] = 0.11[.03], p = 6.51e-05; Figure 3F).

### γ-secretase composite scores predict rate of change in biomarker, clinical and cognitive features in *PSEN1* ADAD carriers

We next assessed how longitudinal rates of change in core biomarker, clinical, and cognitive measures of interest (Figure 4A, C, E, and G) may be related to cell-based measures of γ-secretase function. Lower (more pathogenic) γ-secretase composite measures were associated with faster increase in β-amyloid PET signal (B[SE] = −7.2e-04[2e-04], p = 0.003; Figure 4B), as well as more rapid decreases in hippocampal volume (B[SE] = 4.56[0.7], p = 2.96e-10; Figure 4D), MMSE (B[SE] = 0.02[0.007], p = 3.02e-04; Figure 4F), and logical memory-delayed (B[SE] = 0.006[0.001], p = 2.63e-06; Figure 4H).

**Figure 4.**
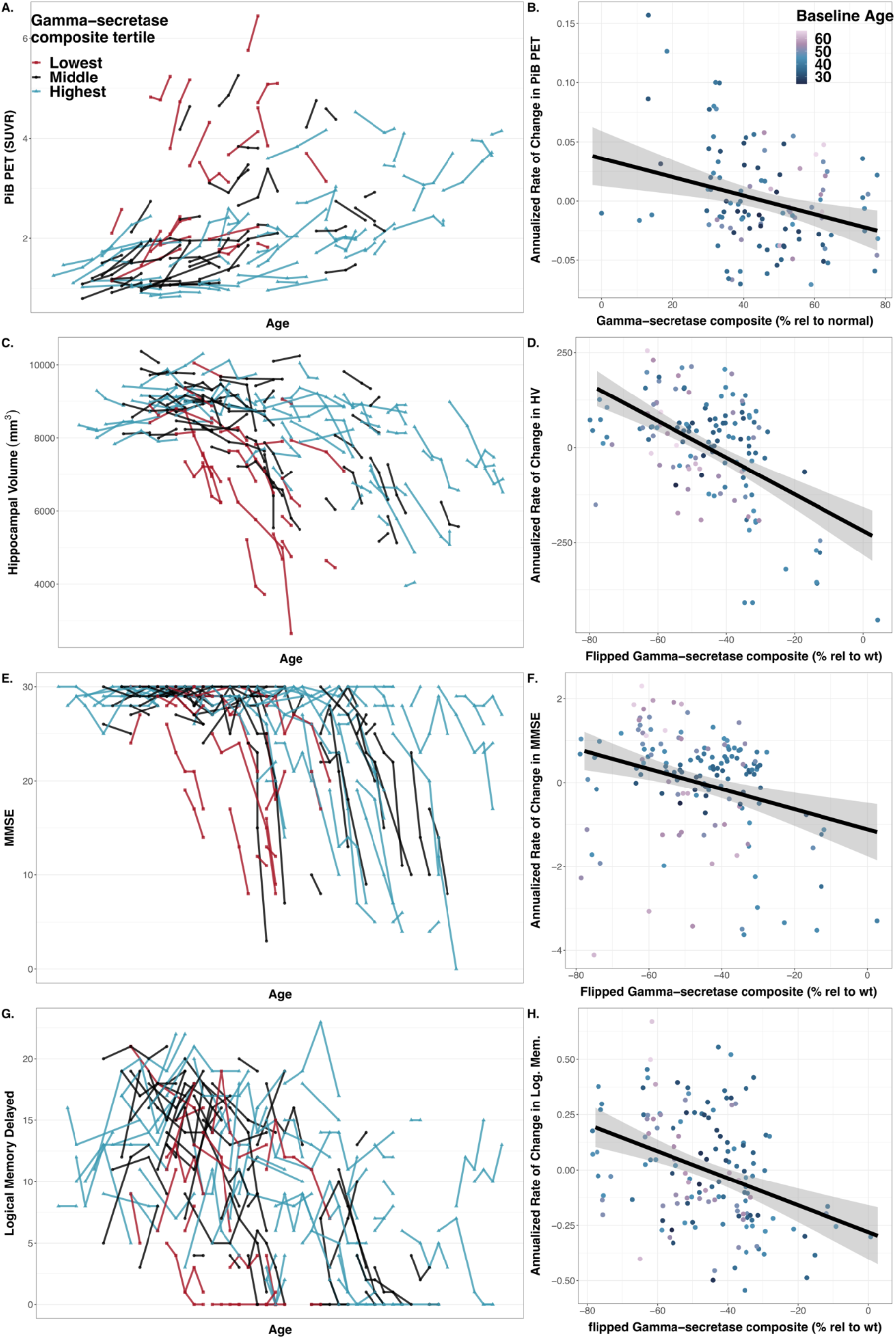
Lower γ-secretase function across *PSEN1* variants is associated with more rapid worsening in biomarker, clinical and cognitive measures in ADAD. Individual longitudinal measures are plotted for PiB-PET SUVR (Aβ burden; **A**), hippocampal volume (HV; mm^3^; **C**), Mini Mental Status Exam (MMSE; **E**), and logical memory delayed (Log. Mem.; **G**). Colors indicate whether an individual’s *PSEN1* variant is in the low (red; most pathogenic tertile), middle (black; intermediate pathogenicity), or high (blue; least pathogenic) tertile for γ-secretase function. To facilitate statistical comparison, age-adjusted, individual-level slopes for each biomarker and cognitive outcomes of interest were extracted from linear mixed effects models and plotted against γ-secretase function (Aβ PET – **B**; Hippocampal Volume – **D**; MMSE – **F**; Logical Memory – **H**). Significant associations were observed between rate-of-change in each measure and variant-level γ-secretase function. Individual data points are jittered to maintain blinding.

### Multimodal models suggest variant level variations in γ-secretase function may impact the clinical, cognitive, and biomarker progression of ADAD

To assess how differences in γ-secretase function may alter the observed clinical, cognitive, and biomarker course of ADAD, we performed a tertile split of variants represented in DIAN and plotted the standardized change in core clinical, cognitive, and biomarker measures across the lifespan for individuals carrying *PSEN1* pathogenic mutations in the lowest, middle, or highest tertile of γ-secretase function (Figure 5A-C). To complement this group-level analysis, we used all available data to visualize putative disease trajectories for carriers of 2 hypothetical variants with 25% and 75% of normal γ-secretase function, respectively (Figure 5D-I). Consistent with the foregoing analyses, these visualizations demonstrate an earlier shift in AAO, clinical and biomarker abnormalities in carriers of variants with lower (more pathogenic) γ-secretase composite scores. A similar pattern was seen when using estimated years to symptom onset (EYO; calculated by subtracting history-derived AAO from the participant’s age) rather than age (Figure S2A-B). This suggests that cell-based assessments of γ-secretase add to the prediction of disease trajectories even after history-derived information about AAO is included in statistical models. However, while a shift in the timing of symptom and biomarker changes was seen across a range of γ-secretase function, it is notable that elevations in pathological Aβ burden (higher Aβ PET or lower CSF Aβ42) and tau (higher CSF phosphorylated-tau181) appear to be early events across all tertiles and precede changes in neurodegenerative markers (hippocampal volume, FDG PET) and clinical measures of impairment (CDR-SB). Together, these results suggest that, while mutation-level variations in γ-secretase function may impact the age of symptom onset and rate of ADAD progression, the ordering of clinical, cognitive, and biomarker changes is not itself fundamentally altered by differences in γ-secretase function.

**Figure 5.**
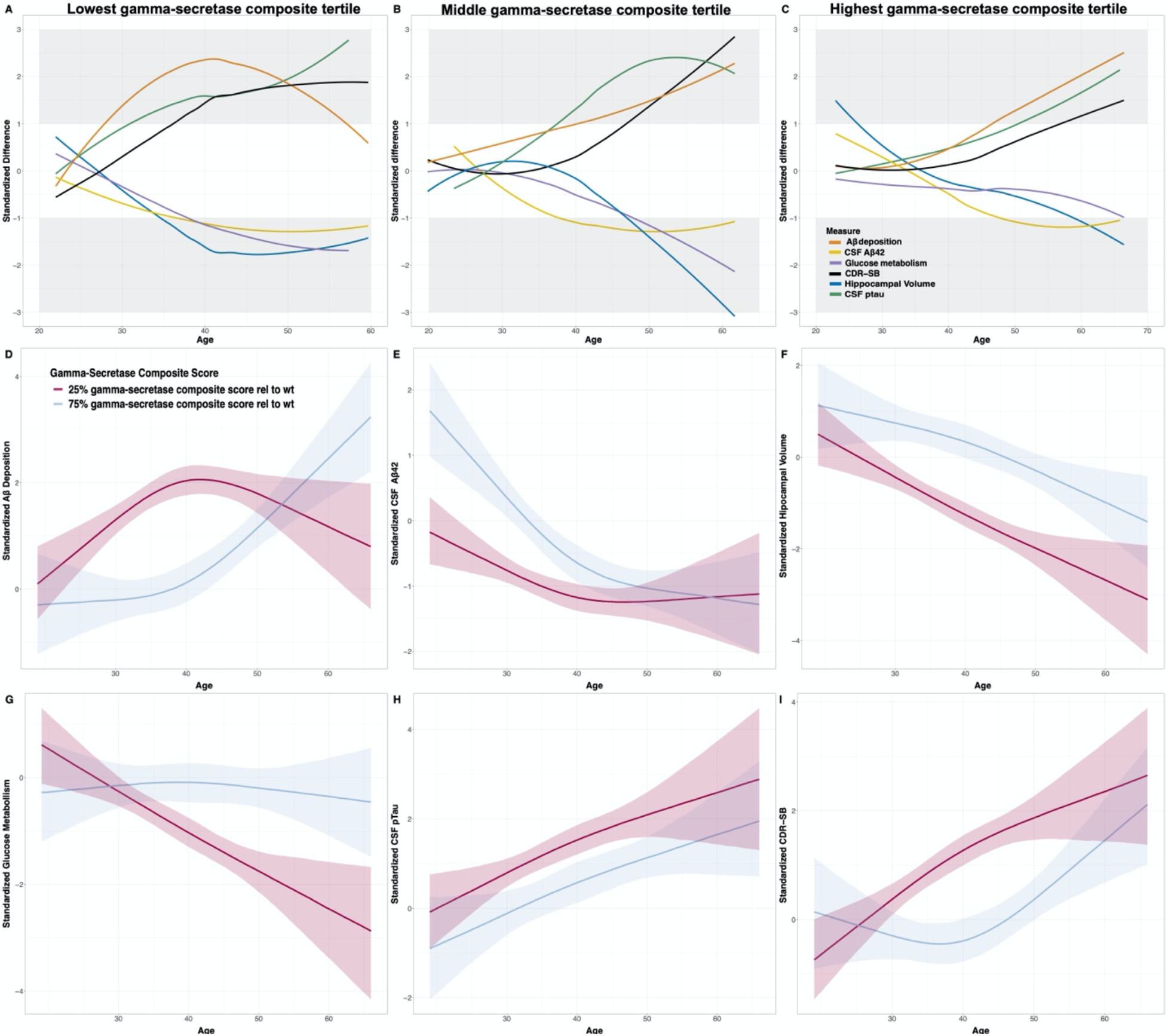
Variant-level variations in γ-secretase function are broadly associated with the clinical and biomarker course of ADAD and can be used to predict ADAD progression for novel *PSEN1* pathogenic variants. To visualize how variant-level differences in γ-secretase function may broadly alter the clinical, cognitive, and biomarker course of ADAD, we plotted the standardized differences in core clinical, cognitive, and biomarker measures across the lifespan for DIAN-Obs participants carrying *PSEN1* pathogenic variants in the lowest (**A**), middle (**B**), or highest (**C**) tertile of γ-secretase function compared to non-carriers. This group-level analysis was followed by an individual-level visualization of disease trajectories for pathogenic variant carriers of 2 possible variants with 25% and 75% of normal γ-secretase function, respectively (D-I).

## DISCUSSION

The study of ADAD forms a cornerstone for understanding AD pathobiology, informing the development of new AD biomarkers, broadly influencing the design of animal models of AD, and advancing therapeutic approaches for both LOAD and ADAD. While ADAD-causing *PSEN1* pathogenic variants are highly penetrant, remarkable heterogeneity in the AAO, rates of disease progression, and biomarker changes is seen across variants. Here we examined whether variant level differences in γ-secretase processing of APP may be critical determinants of the age at which *PSEN1* pathogenic variant carriers manifest cognitive changes and develop abnormalities in core biomarkers for AD. Using a cell-based model system to assess the production of a broad set of Aβ peptides, we observed that mutation level differences in γ-secretase processing of APP strongly predicted the age at which progressive cognitive symptoms are manifest (AAO) for a given variant. In addition to predicting AAO, we observed that a summary measure of γ-secretase function was associated with the rates of functional (CDR-SB) and cognitive decline. This composite measure of γ-secretase function was also associated with all fluid and imaging biomarker measures of AD pathobiology and neurodegeneration we examined, including CSF measures of AD pathology, Aβ PET, FDG-PET, and MRI gray matter volumes. These results suggest that accounting for mutation-level γ-secretase dysfunction may clarify treatment effects in ADAD clinical trials and may also be useful in predicting approximate AAO and disease trajectories for *PSEN1* pathogenic variants that are novel or lack sufficient clinical history. Mechanistically, these results highlight the direct importance of γ-secretase function to AD pathobiology and provide support for γ-secretase modulation as a potentially powerful AD therapeutic and preventative strategy^36^.

Using a human cell-based model system, we found substantial variability in the production of shorter (Aβ-37 and 38) to longer (Aβ-42 and 43) Aβ peptides across *PSEN1* pathogenic variants. The high-throughput nature of our human-cell-based system enabled characterization of a large amount of *PSEN1* variants allowing a more thorough comparison with AAO compared to what has been assessed precviously^26^, which in turn improves the generalizability of our findings with respect to autosomal dominant Alzheimer’s disease. In this same system, we observed that the Aβ42/40 ratio was increased in a majority but not all variants examined. We first examined 106 *PSEN1* variants with available AAO but no available clinical and biomarker data and observed that both the ratio of short-to-long Aβ peptide production 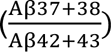 and the more commonly used Aβ 42/40 ratio were predictive of AAO. These results in PSEN1 variants without biomarker, cognitive, and clinical data are concordant with the recent findings in a much small number of *PSEN1* variants from Petit and colleagues^26^, a study which used virally induced mouse embryonic fibroblasts as a model system.

Intriguingly, the short-to-long and Aβ 42/40 ratios were independently associated with AAO, and a composite measure derived from these two ratios provided stronger associations with AAO than either alone. This composite measure was carried forward to examine AD progression in a non-overlapping set of *PSEN1* carriers with available clinical and biomarker measures of AD progression.

We found that the γ-secretase composite measure explained substantial biomarker heterogeneity across *PSEN1* pathogenic mutation carriers using available data from DIAN. Focusing on associations between the γ-secretase composite and Aβ PET, we observed that individuals who carried pathogenic variants in the lowest (most pathogenic) tertile had, on average, 1.4 SUVR units greater Aβ PET signal compared to those with variants in the highest (least pathogenic) tertile, after accounting for age and sex. Putting these differences in the context of the conventionally used threshold for Aβ PET positivity in this cohort and with this Aβ PET tracer (cortical PiB PET SUVR positive at >= 1.42 SUVR^30^), carriers of pathogenic variants in the lowest (most pathogenic) tertile of the γ-secretase composite would, on average, become amyloid positive ∼15 years sooner than those carrying *PSEN1* variants in the highest (least pathogenic) tertile (estimated mean age of 22.6 and 37.2 years for low and high tertiles, respectively). Consistent with cross-sectional results, analysis of longitudinal data demonstrated that Aβ PET signal increased more quickly in individuals carrying *PSEN1* variants with lower (more pathogenic) γ-secretase composite scores compared to those with higher composite scores. Though further work is needed to elucidate regional differences, the results here also suggest that variants with low γ-secretase composite values may have greater striatal Aβ PET signal compared to carriers of *PSEN1* variants with high composite values.

Cerebrospinal fluid measures of AD pathology also had significant associations with the variant-level γ-secretase composite, including CSF measures of Aβ42/40 and phosphorylated-tau181, and phosphorylated-tau217. In addition to associations with core measures of AD pathobiology, γ-secretase composite scores correlated with FDG PET and structural MRI, two commonly used neuroimaging measures of neurodegeneration. Together with the described associations with AAO and functional impairment measures, this broad pattern of association between γ-secretase function with core AD and neurodegenerative markers further support the hypothesis that heterogeneity in ADAD can broadly be understood in the context of mutation-level differences in γ-secretase dysfunction. Importantly, despite variations in AAO and biomarker trajectories across variants, the results here also support prior observations that pathologic changes in Aβ and tau consistently precede neurodegenerative changes and clinical impairment across *PSEN1* variants^25, 26, 37^. Additionally, as the γ-secretase composite used here was derived to predict AAO, it remains quite possible that different formulations of this summary measure of γ-secretase function will be better optimized to predict particular biomarker trajectories in clinical trial settings.

The results here need to be considered in the context of certain limitations. Though AAO information was available on all *PSEN1* variants examined, only a subset had available clinical, cognitive, and biomarker data from the DIAN-Obs study. Accordingly, data from new pathogenic variants in DIAN and in the broader scientific literature will need to be considered. Similarly, tau PET data will need to be considered in future analyses as they become available. Further, it will be important to determine if processing of γ-secretase substrates besides APP is additionally informative for understanding the differential pathogenicity across *PSEN1* variants. Lastly, it will be critical to extend the cell-based model system to allow for the characterization of pathogenic variants in *PSEN2* and *APP*, as ∼20% of ADAD-causing variants are found within these genes.

Despite these limitations, these findings indicate that variant level differences in γ-secretase function are highly associated with core measures of clinical, biomarker, and cognitive, progression of ADAD, and suggest that these variant-level differences explain a large portion of the heterogeneity seen across *PSEN1* pathogenic mutation carriers. As the biochemical measures of γ-secretase function used here are derived outside the context of family polygenic factors, age, and estimated years to symptom onset, they offer a potentially unique channel of information with respect to disease progression compared to these other factors and may thus represent useful complements to these measures in ADAD clinical trials. Similarly, as these cell-based measures are divorced from family polygenic factors and age, they may prove to be valuable in gauging the impact of genetic and lifestyle factors that may confer resilience or risk for ADAD onset and progression. Cell-based measures of γ-secretase function may be particularly useful to assess pathogenicity, potential AAO, and predicted AD trajectories for newly discovered *PSEN1* pathogenic variants or for variants lacking sufficient clinical history. Importantly, these results also strongly support the continued investigation of γ-secretase modulators as an AD therapeutic strategy. Lastly, by demonstrating a tight linkage between the processing of APP by γ-secretase in a cell-based model system and ADAD disease trajectories *in vivo*, the results here provide strong support for the hypothesis that dysregulation of APP processing is central to AD pathogenesis, offing support for some formulations of the “Aβ hypothesis” of AD pathogenesis.

## ACKNOWLEDGMENTS

Data collection and sharing for this project was supported by the Dominantly Inherited Alzheimer Network (DIAN, U19AG032438) funded by the National Institute on Aging (NIA), the Alzheimer’s Association (SG-20-690363-DIAN), the German Center for Neurodegenerative Diseases (DZNE), Raul Carrea Institute for Neurological Research (FLENI). Partial support by the Research and Development Grants for Dementia from Japan Agency for Medical Research and Development, AMED, and the Korea Health Technology R&D Project through Korea Health Industry Development Institute (KHIDI), Spanish Institute of Health Carlos III (ISCIII), Canadian Institutes of Health Research (CIHR), Canadian Consortium of Neurodegeneration and Aging, Brain Canada Foundation and Fonds de Recherche du Québec – Santé. This manuscript has been reviewed by DIAN Study investigators for scientific content and consistency of data interpretation with previous DIAN Study publications. We acknowledge the altruism of the participants and their families, and contributions of the DIAN research and support staff at each of the participating sites for their contributions to this study.

## REFERENCES

1. Lagomarsino VN, Pearse RV, Liu L, et al. Stem cell-derived neurons reflect features of protein networks, neuropathology, and cognitive outcome of their aged human donors. Neuron. 2021;109(21):3402–3420.e9. doi:10.1016/j.neuron.2021.08.003

2. Hsu S, Gordon BA, Hornbeck R, et al. Discovery and validation of autosomal dominant Alzheimer’s disease mutations. Alz Res Therapy. 2018;10(1):67. doi:10.1186/s13195-018-0392-9

3. Karch CM, Hernández D, Wang JC, et al. Human fibroblast and stem cell resource from the Dominantly Inherited Alzheimer Network. Alzheimers Res Ther. 2018;10(1):69. doi:10.1186/s13195-018-0400-0

4. Lanoiselée HM, Nicolas G, Wallon D, et al. APP, PSEN1, and PSEN2 mutations in early-onset Alzheimer disease: A genetic screening study of familial and sporadic cases. PLoS Med. 2017;14(3):e1002270. doi:10.1371/journal.pmed.1002270

5. De Strooper B, Iwatsubo T, Wolfe MS. Presenilins and γ-secretase: structure, function, and role in Alzheimer Disease. Cold Spring Harb Perspect Med. 2012;2(1):a006304. doi:10.1101/cshperspect.a006304

6. Fernandez MA, Klutkowski JA, Freret T, Wolfe MS. Alzheimer presenilin-1 mutations dramatically reduce trimming of long amyloid β-peptides (Aβ) by γ-secretase to increase 42- to-40-residue Aβ. J Biol Chem. 2014;289(45):31043–31052. doi:10.1074/jbc.M114.581165

7. Liu L, Lauro BM, Wolfe MS, Selkoe DJ. Hydrophilic loop 1 of Presenilin-1 and the APP GxxxG transmembrane motif regulate γ-secretase function in generating Alzheimer-causing Aβ peptides. Journal of Biological Chemistry. 2021;296:100393. doi:10.1016/j.jbc.2021.100393

8. Wolfe MS. Dysfunctional γ-Secretase in Familial Alzheimer’s Disease. Neurochem Res. 2019;44(1):5–11. doi:10.1007/s11064-018-2511-1

9. Wolfe MS, Selkoe DJ. Giving Alzheimer’s the old one-two. Cell. 2010;142(2):194–196. doi:10.1016/j.cell.2010.07.006

10. Szaruga M, Munteanu B, Lismont S, et al. Alzheimer’s-Causing Mutations Shift Aβ Length by Destabilizing γ-Secretase-Aβn Interactions. Cell. 2017;170(3):443–456.e14. doi:10.1016/j.cell.2017.07.004

11. Wolfe MS. Probing Mechanisms and Therapeutic Potential of γ-Secretase in Alzheimer’s Disease. Molecules. 2021;26(2):E388. doi:10.3390/molecules26020388

12. Nie P, Vartak A, Li YM. γ-Secretase inhibitors and modulators: Mechanistic insights into the function and regulation of γ-Secretase. Semin Cell Dev Biol. 2020;105:43–53. doi:10.1016/j.semcdb.2020.03.002

13. Raven F, Ward JF, Zoltowska KM, et al. Soluble Gamma-secretase Modulators Attenuate Alzheimer’s β-amyloid Pathology and Induce Conformational Changes in Presenilin 1. EBioMedicine. 2017;24:93–101. doi:10.1016/j.ebiom.2017.08.028

14. Psen-1| Alzforum. (n.d.). Retrieved August 09, 2021, from https://www.Alzforum.Org/Mutations/Psen-1.

15. Ryan NS, Nicholas JM, Weston PSJ, et al. Clinical phenotype and genetic associations in autosomal dominant familial Alzheimer’s disease: a case series. The Lancet Neurology. 2016;15(13):1326–1335. doi:10.1016/S1474-4422(16)30193-4

16. Ryman DC, Acosta-Baena N, Aisen PS, et al. Symptom onset in autosomal dominant Alzheimer disease: a systematic review and meta-analysis. Neurology. 2014;83(3):253–260.

17. Lippa CF, Swearer JM, Kane KJ, et al. Familial Alzheimer’s disease: site of mutation influences clinical phenotype. Ann Neurol. 2000;48(3):376–379.

18. Wegiel J, Wisniewski HM, Kuchna I, et al. Cell-Type-Specific Enhancement of Amyloid-β Deposition in a Novel Presenilin-1 Mutation (P117L): Journal of Neuropathology and Experimental Neurology. 1998;57(9):831–838. doi:10.1097/00005072-199809000-00004

19. Klunk WE, Price JC, Mathis CA, et al. Amyloid Deposition Begins in the Striatum of Presenilin-1 Mutation Carriers from Two Unrelated Pedigrees. Journal of Neuroscience. 2007;27(23):6174–6184. doi:10.1523/JNEUROSCI.0730-07.2007

20. Tang M, Ryman DC, McDade E, et al. Neurological manifestations of autosomal dominant familial Alzheimer’s disease: a comparison of the published literature with the Dominantly Inherited Alzheimer Network observational study (DIAN-OBS). The Lancet Neurology. 2016;15(13):1317–1325.

21. Ryan NS, Rossor MN. Correlating familial Alzheimer’s disease gene mutations with clinical phenotype. Biomarkers Med. 2010;4(1):99–112. doi:10.2217/bmm.09.92

22. Chhatwal JP, Schultz SA, McDade E, et al. Variant-dependent heterogeneity in amyloid β burden in autosomal dominant Alzheimer’s disease: cross-sectional and longitudinal analyses of an observational study. Lancet Neurol. 2022;21(2):140–152. doi:10.1016/S1474-4422(21)00375-6

23. Kumar-Singh S, Theuns J, Van Broeck B, et al. Mean age-of-onset of familial alzheimer disease caused by presenilin mutations correlates with both increased Aβ42 and decreased Aβ40. Hum Mutat. 2006;27(7):686–695. doi:10.1002/humu.20336

24. Szaruga M, Veugelen S, Benurwar M, et al. Qualitative changes in human γ-secretase underlie familial Alzheimer’s disease. J Exp Med. 2015;212(12):2003–2013. doi:10.1084/jem.20150892

25. Liu L, Lauro BM, He A, et al. Identification of the Aβ37/42 peptide ratio in CSF as an improved Aβ biomarker for Alzheimer’s disease. Alzheimers Dement. Published online March 12, 2022. doi:10.1002/alz.12646

26. Petit D, Fernández SG, Zoltowska KM, et al. Aβ profiles generated by Alzheimer’s disease causing PSEN1 variants determine the pathogenicity of the mutation and predict age at disease onset. Mol Psychiatry. 2022;27(6):2821–2832. doi:10.1038/s41380-022-01518-6

27. Bateman RJ, Xiong C, Benzinger TLS, et al. Clinical and Biomarker Changes in Dominantly Inherited Alzheimer’s Disease. New England Journal of Medicine. 2012;367(9):795–804. doi:10.1056/NEJMoa1202753

28. Wechsler D. Wechsler Memory Scale (3rd ed.): Administration and scoring manual. Published online 1997.

29. Barthélemy NR, Li Y, Joseph-Mathurin N, et al. A soluble phosphorylated tau signature links tau, amyloid and the evolution of stages of dominantly inherited Alzheimer’s disease. Nat Med. 2020;26(3):398–407. doi:10.1038/s41591-020-0781-z

30. Su Y, D’Angelo GM, Vlassenko AG, et al. Quantitative Analysis of PiB-PET with FreeSurfer ROIs. PLOS ONE. 2013;8(11):e73377. doi:10.1371/journal.pone.0073377

31. Su Y, Blazey TM, Owen CJ, et al. Quantitative Amyloid Imaging in Autosomal Dominant Alzheimer’s Disease: Results from the DIAN Study Group. PLoS One. 2016;11(3):e0152082. doi:10.1371/journal.pone.0152082

32. Gordon BA, Blazey TM, Su Y, et al. Spatial patterns of neuroimaging biomarker change in individuals from families with autosomal dominant Alzheimer’s disease: a longitudinal study. The Lancet Neurology. 2018;17(3):241–250.

33. McKay NS, Gordon BA, Hornbeck RC, et al. Neuroimaging within the Dominantly Inherited Alzheimer’s Network (DIAN): PET and MRI. Neuroscience; 2022. doi:10.1101/2022.03.25.485799

34. Gu L, Guo Z. Alzheimer’s Aβ42 and Aβ40 peptides form interlaced amyloid fibrils. J Neurochem. 2013;126(3):305–311. doi:10.1111/jnc.12202

35. Cullen N, Janelidze S, Palmqvist S, et al. Association of CSF Aβ38 Levels With Risk of Alzheimer Disease-Related Decline. Neurology. 2022;98(9):e958–e967. doi:10.1212/WNL.0000000000013228

36. Toyn JH, Boy KM, Raybon J, et al. Robust Translation of -Secretase Modulator Pharmacology across Preclinical Species and Human Subjects. Journal of Pharmacology and Experimental Therapeutics. 2016;358(1):125–137. doi:10.1124/jpet.116.232249

37. Sun L, Zhou R, Yang G, Shi Y. Analysis of 138 pathogenic mutations in presenilin-1 on the in vitro production of Aβ42 and Aβ40 peptides by γ-secretase. Proc Natl Acad Sci U S A. 2017;114(4):E476–E485. doi:10.1073/pnas.1618657114

