## Supplemental Materials for "Functional variations in gamma-secretase activity are critical determinants of the clinical, biomarker, and cognitive progression of autosomal dominant Alzheimer’s disease"

**Supplemental Methods:**

*Generation of Presenilin1/2 double knockout cell line (PSEN1/2 dKO):*

Homozygous human PSEN1/2 dKO HEK-293 cell lines were generated using a clustered regularly interspaced short palindromic repeat (CRISPR)/Cas9 nuclease-mediated system (Addgene). The guide RNAs were designed using the CRISPR Design software package (http://crispr.mit.edu/) to minimize potential off-target effects. Two oligo pairs for Presenilin 1 (PSEN1) and three oligo pairs for Presenilin 2 (PSEN2) were cloned into vector PX459 (Addgene, #62988) to express five gRNAs targeting PSEN1/2. After transfection of PX459 into HEK293 cells for 24 h, puromycin selection was undertaken for 1 week. PSEN1/2 dKO cells were isolated via limiting dilution cloning and confirmed by western blots.

*Generation of presenilin-1 and C99 expression vectors:*

PcDNA 3.1 vector harboring the wild-type human presenilin-1 and PcDNA 3.1 vector harboring the wild-type (wt) human APP-C99 with signal peptide 1–16 were used for the template to generate presenilin-1 and C99 expression vectors. To introduce mutations, the template was amplified by PCR into two DNA fragments with an overlapping sequence containing the mutated location. Overlap PCR was done to generate the whole open reading frame containing the mutation. PCR products were subcloned into the parental vector. Vectors were sequenced from both 5' and 3' ends to confirm successful mutagenesis. *PSEN1* constructs for all 55 FAD mutations represented in the Dominantly Inherited Alzheimer's Network (DIAN) observational study (data freeze version 12; list of FAD mutations included in Table S1) were generated and transiently coexpressed in HEK-293 *PSEN1*/2 dKO line together with wt human *APP* (Figure 1A).

*Tissue culture and transfection of adherent cells:*

Adherent HEK cells were cultured in complete growth media: Dulbecco's Modified Eagle's Medium (DMEM) supplemented with 10% fetal bovine serum (FBS), 2 mM Lglutamine, 10 units/ml penicillin, and 10 mg/ml streptomycin. For transfection, adherent HEK cells were seeded in 24-well dishes at a density of 5 x 105 cells per well. Transfection was carried out with jetPrime reagent. Cells were incubated for 24 h and media were changed for conditioning after another 12 h, at which time the conditioned media were harvested for ELISA, and the cells were harvested for western blots.

*Aβ ELISA:*

Conditioned media from transfected HEK cells were harvested and diluted with 1% BSA in wash buffer (TBS supplemented with 0.05% Tween). For Aβx-37, x-38, x-40, x-42, and x-43 assays, each well of an uncoated 96-well multiarray plate (Meso Scale Discovery, #L15XA-3) was coated with 30 uL of a PBS solution containing 3 µg/ml of 266 capture antibody (Elan) and incubated at room temperature overnight. A detection antibody solution was prepared with biotinylated monoclonal antibody recognizing the respective C-terminal residue of each Aβ peptide, plus 100 ng/ml Streptavidin SulfoTAG (Meso Scale Discovery, #R32AD-5) and 1% BSA diluted in wash buffer. Following overnight incubation, 50 µl/well of the CM sample and 25 µl/well of the detection antibody solution were incubated for 2 h at room temperature with shaking at >300 rpm, washing wells with wash buffer between incubations. The plate was read and analyzed according to manufacturer's protocol. Aβ ELISAs were performed in triplicate and values were averaged.

**Imaging Analyses:**

*MRI*

DIAN Imaging data was screened for protocol compliance and artifacts. All sites used a 3T scanner that was qualified for use at study initiation and was required to pass regular quality control assessments. Volumetric T1-weighted images were acquired for all participants and were processed using FreeSurfer v 5.3 (http://surfer.nmr.mgh.harvard.edu/) [1,2] and the Desikan-Killany atlas to produce regional estimates of grey matter volume within brain regions. Our analyses focused on the hippocampus as the *a priori* region of interest (ROI). Hippocampal volumes were averaged across left and right hemispheres and adjusted for total intracranial volume prior to statistical analysis.

*PET*

Amyloid imaging was performed with a bolus injection of ~15 mCi of [$^{11}$C] PiB. Dynamic acquisition consisted of either a 70-min scan starting at injection or a 30-min scan beginning 40 min post injection. For analysis, the PiB PET data in the common time frame between 40–70 min was used. Metabolic imaging with [$^{18}$F] FDG-PET was performed with a 30-min 3D dynamic acquisition beginning 30 min after injection. The last 25 min of each FDG scan was used for analysis purposes. The ADNI PET Core verified that all PET images were acquired using the established protocol and substantially free of artifacts. Using FreeSurfer ROIs, standardized uptake value ratios (SUVRs) were calculated using the cerebellar grey matter as a reference region (PET Unified Pipeline, https://github.com/ysu001/PUP). To minimize the impact of partial volume effects on the PET signal, an RSF-based approach for partial volume correction was used for all regional PET measurements.[3] MRI and PET data acquisition and processing has been described in detail in previous studies [4,5]. As previously described, a composite measure for mean cortical Aβ deposition measure was generated using the average across the left and right lateral orbitofrontal, medial orbitofrontal, rostral middle frontal, superior frontal, superior temporal, middle temporal, and precuneus regions [6,7]. We chose a single precuneus (average of both hemispheres) region for FDG based on prior work in this and other ADAD cohorts indicating this region as the earliest effected by a number of different imaging measures. Exploratory regional analyses examined additional regions (see Figure 2D-F) for which FreeSurfer data was available.

**CSF Analyses:**

CSF was obtained using procedures consistent with the biofluid protocol of the Alzheimer's Disease Neuroimaging Initiative (ADNI). Briefly, CSF was drawn using 21-22g Sprotte or Quincke spinal needles into polypropylene tubes, followed by placement on dry ice and shipment to the DIAN Biomarker Core at Washington University. Frozen samples were then thawed, aliquoted, and stored at -84 degrees C until assayed. CSF assays for Aβ 40, and Aβ 42, total tau, and phospho-tau 181 were performed at the DIAN Biomarker Core using an automated immunoassay system (LUMIPULSE G1200,

Fujirebio, Malvern, PA) according to manufacturer's specifications. CSF phospho-tau 181 was log-transformed prior to analyses.

As previously described [8], thawed CSF samples were additionally analyzed by nano liquid chromatography coupled to high-resolution tandem mass spectrometry (HRMS/MS) using parallel reaction monitoring and HCD fragmentation. A ratio of phosphorylation on T217 were measured using the ratio of the HRMS/MS transitions from phosphorylated peptides and the corresponding non-phosphorylated peptides. Each phosphorylated/non-phosphorylated peptide endogenous ratio was normalized using the ratio measured on the HRMS/MS transitions of the corresponding phosphorylated/non-phosphorylated peptide internal standards.

**Harmonization of data from Liu et al., 2022:**

A total of 131 variants were characterized in Liu et al., 2022[9] using the same model system and immunoassays used here to characterize the 55 variants from DIAN. To facilitate comparisons, the γ-secretase composite scores were adjusted across the two studies using data from 24 *PSEN1* pathogenic variants that were examined in both studies. Data from these 24 overlapping variants were used for normalization purposes and were not included in analyses examining associations between the γ-secretase composite from the 107 non-overlapping variants described in Liu et al., 2022 data set and AAO (Figure 1).

**SUPPLEMENTARY TABLES AND FIGURES:**

**Table S1. Aβ secretion ratios from *PSEN1* dKO 293HEK cells co-transfected with normal or mutant *PSEN1* and normal APP**

| | $AAO_{gsc}$ | $\frac{A\beta 42}{A\beta 40}$ | Aβ short-to-long ratio | γ-secretase composite (% relative to normal) |
|---|---|---|---|---|
| normal | | 0.09 | 1.22 | 100.00 |
| Ala79Val | 53.71 | 0.16 | 0.60 | 62.45 |
| Met84Val | 54.65 | 0.18 | 0.71 | 64.79 |
| Cys92Ser | 48.44 | 0.16 | 0.32 | 49.23 |
| Phe105Leu | 50.54 | 0.17 | 0.45 | 54.50 |
| Phe105Ser | 50.55 | 0.15 | 0.41 | 54.53 |
| Tyr115His | 41.77 | 0.30 | 0.21 | 32.51 |
| Asn135Tyr | 27.74 | 1.29 | 0.05 | < 1 |
| Asn135Ser | 33.42 | 0.66 | 0.08 | 11.58 |
| Met139Ile | 41.35 | 0.31 | 0.19 | 31.45 |
| Ile143Thr | 37.81 | 0.49 | 0.19 | 22.58 |
| Met146Leu | 41.13 | 0.31 | 0.19 | 30.90 |
| Met146Ile | 43.89 | 0.23 | 0.21 | 37.81 |
| Met146Val | 44.60 | 0.23 | 0.26 | 39.61 |
| Thr147Ile | 34.87 | 0.76 | 0.22 | 15.22 |
| His163Arg | 45.99 | 0.20 | 0.27 | 43.08 |
| Ser169Leu | 34.85 | 0.69 | 0.18 | 15.16 |
| Ser170Phe | 37.14 | 0.39 | 0.06 | 20.90 |
| Leu171Pro | 41.64 | 0.28 | 0.17 | 32.18 |
| Phe176Val | 51.81 | 0.15 | 0.46 | 57.68 |
| Ser178Pro | 41.77 | 0.26 | 0.14 | 32.49 |
| Glu184Asp | 41.53 | 0.28 | 0.16 | 31.91 |
| Ile202Phe | 49.16 | 0.18 | 0.39 | 51.03 |
| Gly206Ala | 53.29 | 0.18 | 0.64 | 61.38 |
| Gly209Val | 33.27 | 0.66 | 0.07 | 11.20 |
| Gly209Glu | 42.36 | 0.31 | 0.25 | 33.99 |
| Ser212Tyr | 46.69 | 0.21 | 0.33 | 44.85 |
| Gly217Arg | 40.63 | 0.27 | 0.10 | 29.65 |
| Leu219Pro | 44.33 | 0.20 | 0.17 | 38.92 |
| Gln222His | 41.81 | 0.26 | 0.15 | 32.61 |

| | | | | |
|---|---|---|---|---|
| Leu226Arg | 40.85 | 0.28 | 0.12 | 30.20 |
| Ile229Phe | 40.56 | 0.49 | 0.35 | 29.47 |
| Ser230Asn | 40.90 | 0.31 | 0.17 | 30.33 |
| Met233Thr | 40.28 | 1.15 | 0.71 | 28.77 |
| Met233Leu | 41.86 | 0.31 | 0.22 | 32.74 |
| Leu235Val | 50.75 | 0.17 | 0.48 | 55.03 |
| Ile238Met | 48.25 | 0.22 | 0.45 | 48.76 |
| Thr245Pro | 34.63 | 0.55 | 0.07 | 14.59 |
| Ala260Gly | 43.34 | 0.32 | 0.33 | 36.45 |
| Ala260Val | 44.82 | 0.18 | 0.15 | 40.15 |
| Val261Phe | 54.38 | 0.14 | 0.58 | 64.14 |
| Pro264Leu | 43.42 | 0.24 | 0.20 | 36.64 |
| Pro267Leu | 48.61 | 0.17 | 0.35 | 49.66 |
| Arg269His | 50.08 | 0.18 | 0.45 | 53.34 |
| Leu271Val | 44.67 | 0.25 | 0.29 | 39.77 |
| Ala275Val | 48.02 | 0.22 | 0.43 | 48.17 |
| Glu280Ala | 42.08 | 0.29 | 0.21 | 33.28 |
| Glu280Gly | 44.13 | 0.20 | 0.16 | 38.42 |
| Phe283Leu | 46.92 | 0.17 | 0.25 | 45.43 |
| Tyr288His | 35.51 | 0.49 | 0.07 | 16.82 |
| E9 Del | 43.51 | 0.24 | 0.21 | 36.87 |
| Ser290Cys | 59.13 | 0.10 | 0.69 | 76.03 |
| Cys410Tyr | 41.88 | 0.27 | 0.17 | 32.79 |
| Ala426Pro | 49.89 | 0.18 | 0.45 | 52.86 |
| Ala431Glu | 47.48 | 0.14 | 0.20 | 46.82 |
| Ile439Val | 59.77 | 0.10 | 0.73 | 77.65 |

**$AAO_{gsc}$** = $\gamma$-secretase composite derived AAO

**Table S2. Cross-sectional study sample characteristics for DIAN-Obs participants.**

| Characteristic | Value |
|---|---|
| Age, years | 40.0 (10.3) |
| EYO, years | -5.8 (10.4) |
| AAO, years | 46.8 (7.8) |
| APOE4+, % | 29.3 |
| Female, % | 58.0 |
| Education, years | 14.4 (2.9) |
| CDR (0, 0.5, 1+), % | 58, 22.9, 19.1 |

Mean (SD) values presented unless otherwise noted.

EYO = Estimated years to symptom onset; AAO= Estimated age at symptom onset; CDR = Global Clinical Dementia Rating Scale

**Table S3. Longitudinal study sample characteristics for DIAN-Obs participants.**

| Characteristic | Value |
|---|---|
| Baseline Age, years | 38.0 (10.4) |
| Baseline EYO, years | -7.5 (10.9) |
| AAO, years | 46.4 (7.7) |
| APOE4+, % | 30.5 |
| Female, % | 57.6 |
| Education, years | 14.4 (2.7) |
| Baseline CDR (0, 0.5, 1+), % | 55.1, 25.4, 19.5 |
| Average years of follow-up | 3.3 (2.1) |

Mean (SD) values presented unless otherwise noted.

EYO = Estimated years to symptom onset; AAO= Estimated age at symptom onset; CDR = Global Clinical Dementia Rating Scale

**Table S4. Aβ ratios uniquely contribute to prediction of age of symptom onset.**

| Term | B (SE) | P-value |
|---|---|---|
| Intercept | 14.91 (4.0) | 3.06e-04 |
| $\frac{\text{short A}\beta 37+38}{\text{long A}\beta 42+43}$ % rel normal | 0.03 (0.01) | 9.15e-05 |
| $\log_{10}(\frac{\text{A}\beta 42}{\text{A}\beta 40}$ % rel normal) | 0.17 (0.04) | 9.30e-05 |

**Figure S1. Schematic showing cell-based methods.**

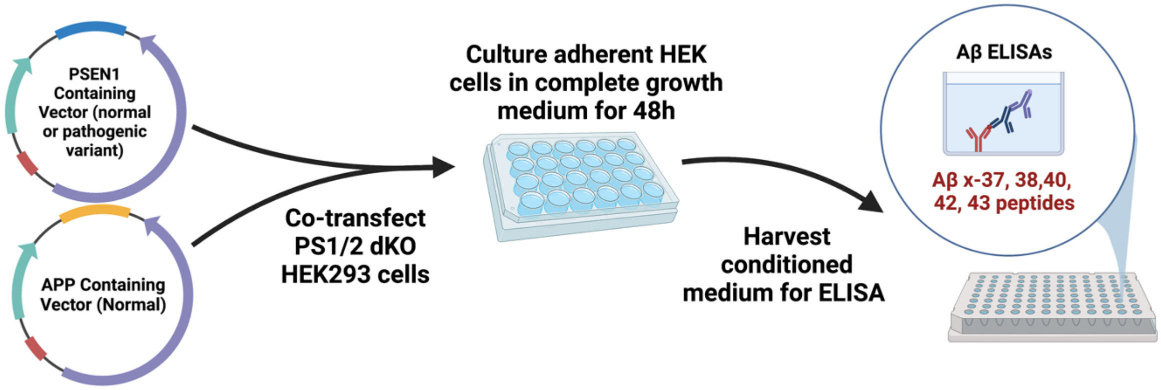

**Figure S2. Variant-level variations in γ-secretase function can be used to predict ADAD progression for novel *PSEN1* pathogenic variants.**

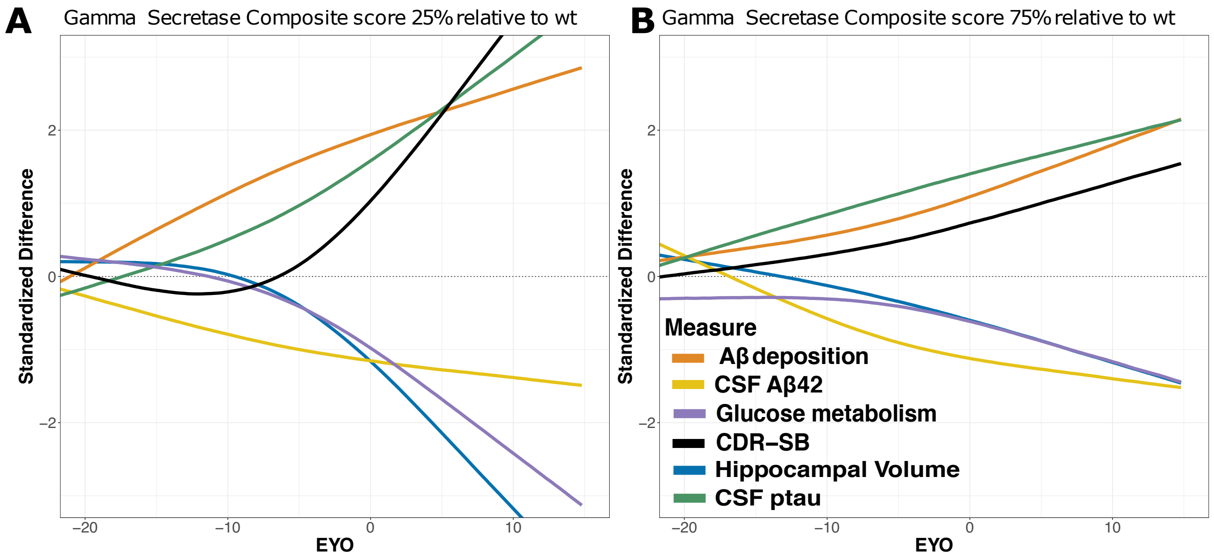


To visualize how variant-level differences in γ-secretase function may broadly alter the clinical, cognitive, and biomarker course of ADAD, an individual-level visualization of disease trajectories for pathogenic variant carriers of 2 possible variants at the 25% (**A**) and 75% (**B**) of normal γ-secretase function are plotted across estimated years to symptom onset (EYO).

### REFERENCES


1. Fischl, B. FreeSurfer. *NeuroImage* **62**, 774–781 (2012).

2. Fischl, B. *et al.* Automatically parcellating the human cerebral cortex. *Cereb Cortex* vol. 14 11–22 (2004).

3. Su, Y. *et al.* Partial volume correction in quantitative amyloid imaging. *Neuroimage* vol. 107 55–64 (2015).

4. Bateman, R. J. *et al.* Clinical and Biomarker Changes in Dominantly Inherited Alzheimer's Disease. *New England Journal of Medicine* **367**, 795–804 (2012).

5. Benzinger, T. L. *et al.* Regional variability of imaging biomarkers in autosomal dominant Alzheimer's disease. *Proceedings of the National Academy of Sciences* vol. 110 E4502–E4509 (2013).

6. Su, Y. *et al.* Quantitative Analysis of PiB-PET with FreeSurfer ROIs. *PLOS ONE* **8**, e73377 (2013).

7. Su, Y. *et al.* Quantitative Amyloid Imaging in Autosomal Dominant Alzheimer's Disease: Results from the DIAN Study Group. *PLoS One* vol. 11 e0152082 (2016).

8. Barthélemy, N. R. *et al.* A soluble phosphorylated tau signature links tau, amyloid and the evolution of stages of dominantly inherited Alzheimer's disease. *Nat Med* **26**, 398–407 (2020).

9. Liu, L. *et al.* Identification of the Aβ37/42 peptide ratio in CSF as an improved Aβ biomarker for Alzheimer's disease. *Alzheimers Dement* (2022) doi:10.1002/alz.12646.
